# Rapid profiling of the preterm infant gut microbiota using nanopore sequencing aids pathogen diagnostics

**DOI:** 10.1101/180406

**Authors:** Richard M. Leggett, Cristina Alcon-Giner, Darren Heavens, Shabhonam Caim, Thomas C. Brook, Magdalena Kujawska, Samuel Martin, Lesley Hoyles, Paul Clarke, Lindsay J. Hall, Matthew D. Clark

## Abstract

The Oxford Nanopore MinION sequencing platform offers near real time analysis of DNA reads as they are generated, which makes the device attractive for in-field or clinical deployment, e.g. rapid diagnostics. We used the MinION platform for shotgun metagenomic sequencing and analysis of gut-associated microbial communities; firstly, we used a 20-species human microbiota mock community to demonstrate how Nanopore metagenomic sequence data can be reliably and rapidly classified. Secondly, we profiled faecal microbiomes from preterm infants at increased risk of necrotising enterocolitis and sepsis. In single patient time course, we captured the diversity of the immature gut microbiota and observed how its complexity changes over time in response to interventions, i.e. probiotic, antibiotics and episodes of suspected sepsis. Finally, we performed ‘real-time’ runs from sample to analysis using faecal samples of critically ill infants and of healthy infants receiving probiotic supplementation. Real-time analysis was facilitated by our new NanoOK RT software package which analysed sequences as they were generated. We reliably identified potentially pathogenic taxa (i.e. *Klebsiella pneumoniae and Enterobacter cloacae*) and their corresponding antimicrobial resistance (AMR) gene profiles within as little as one hour of sequencing. Antibiotic treatment decisions may be rapidly modified in response to these AMR profiles, which we validated using pathogen isolation, whole genome sequencing and antibiotic susceptibility testing. Our results demonstrate that our pipeline can process clinical samples to a rich dataset able to inform tailored patient antimicrobial treatment in less than 5 hours.

## INTRODUCTION

Next generation sequencing (NGS) has revolutionised profiling of environmental and clinical microbial communities. In particular, the culture-independent, sensitive, data-rich nature of metagenomic sequencing, combined with powerful bioinformatics tools, have allowed researchers to start to differentiate patient groups from healthy individuals based on their microbial profiles^1–5^. Microbiota linked health effects include the ability to support immune system development, facilitate dietary metabolism, modulate the metabolome and provide antimicrobial protection^6–8^. Disturbances in the microbiota by, for example, external influences such as antibiotics, have been associated with an increased risk of diseases including ulcerative colitis^9^, obesity, and autoimmune conditions^10, 11^, and within an infectious disease context increased risk of pathogen overgrowth^12^.

NGS platforms are often large capital investments with a considerable physical footprint and generate huge quantities of data using a pipeline which combines many samples into a single sequencing run, which can take days or weeks to run and analyse. These are not ideal attributes for clinical settings. In contrast, the MinION platform introduced by Oxford Nanopore Technologies in 2014 is a pocket-sized sequencing device powered from a laptop’s USB port capable of producing gigabase yields of long reads in real time (see recent review of MinION technology and applications^13^). This low cost, $1000 platform could potentially allow clinical scientists a readily portable and easy-to-use NGS platform that shows great potential for rapid pathogen screening, diagnosis, and treatment strategy design. Crucially, the device operates in ‘real-time’ allowing sequenced reads to be analysed immediately after they are generated. However, due to the ‘real-time’ nature of the platform and its different error profile, it is critical to develop new bioinformatics pipelines to take full advantage of the data, particularly in the clinical arena.

The huge rise in antimicrobial resistance (AMR)^14–16^ highlights the need for new technologies able to identify at-risk individuals, diagnose infectious agents and suggest optimised treatments. The MinION may represent such a technology, and one at-risk patient cohort is premature infants, defined as born before the start of the 37th week of pregnancy, and accounting for 1 in 10 live births globally, and increasing^17^. These infants are born with underdeveloped gut physiology and immunity, which is associated with an increased risk of life threatening infections^18^. Furthermore, preterm infants are often born via Caesarean-section and receive multiple antibiotic courses during their hospitalisation that deplete their gut microbiota, including the beneficial genus *Bifidobacterium*, that may correlate with overgrowth of pathogens linked to diseases such as sepsis and necrotising enterocolitis (NEC)^19, 20^. Notably, these diseases are difficult to diagnose in the early stages and are often associated with sudden serious deterioration. The most common pathogens linked to sepsis and NEC include *Clostridium perfringens, group B streptococcus, Escherichia coli*^21^, *Enterobacter* spp., and *Klebsiella pneumoniae*^22^. Therefore a good diagnostic method must be able to confidently identify (i) microbes to the species level for accurate diagnosis, (ii) species abundance within the microbiota (as these bacteria can be present within the wider community, but not cause disease when at low levels) and (iii) AMR gene repertoires.

Despite technical challenges in metagenomic profiling for diagnostics^23^, MinIONs have been successfully used in medical research on low complexity samples. To date this includes surveillance of the Ebola outbreak in West Africa (using RT-PCR amplicons)^24^, characterisation of bacterial isolates^25^ and DNA spiked into patient samples at known levels^26^ or from samples heavily (~90%) infected with a single species^27, 28^. Diagnostics in metagenomic samples is still challenging due to lower MinION sequence yields and accuracy but is essential as most clinical samples are complex. A recent a study using mock environmental samples correctly assigned taxonomy in single bacterial species in low complexity synthetic communities using older flow cells R7.3^29^, however to date no studies have explored MinION technology in clinical metagenomic samples. Here we demonstrate MinION based metagenomics, first in a controlled (i.e. mock community) setting and then with clinical samples including in ‘real time’. These studies allowed us to determine longitudinal microbiota profiles, gut-associated pathogens linked with sepsis or NEC and their AMR profiles. We benchmarked these data against conventional Illumina sequencing, whole-genome sequencing (WGS) of pathogen isolates and microbiology antibiotic susceptibility testing.

## RESULTS

### Accurate classification of a microbial mock community using MinION sequencing

We benchmarked the efficacy of MinION technology by profiling a bacterial mock community of a staggered abundance mixture (HM-277D, BEI Resources). Sequencing was carried out on R7.3 flow cells using a “2D” library protocol that involves sequencing both strands of DNA. One flow cell produced 148,441 total reads, with 71,675 reads passing default quality filters and a mean read length of 3,047 bp (read size is largely driven by the size of input DNA molecules) with the longest read of 40,561 bp (**Table 1**). Reads were analysed with NanoOK^30^ and produced alignments to the 20 microbial reference sequences with 82-89% identity (consistent with previous published analysis^31^) including long error-free sequence stretches of up to 223 bases (the full NanoOK report is available at https://github.com/richardmleggett/bambi). Coverage of each genome ranged from almost 0× (8 reads of *Actinomyces odontolyticus*) to 13× (7,695 reads of *Streptococcus mutans*), which is consistent with their mock concentrations (**Supplementary table 1**). We benchmarked to ‘conventional’ (Illumina based) metagenomic sequencing using 1 million reads (equivalent to a MiSeq nano flow cell). To simulate an unknown community, reads were BLAST searched against the NCBI nt database and then classified taxonomically using MEGAN6^32^. This showed broadly similar abundance levels across both platforms, but in some cases, MEGAN’s Lowest Common Ancestor algorithm was able to assign a greater proportion of Nanopore reads to species level rather than genus or family, while in other cases, a greater proportion of Illumina reads were able to be assigned to species (**Fig. 1a**). This apparent contradiction is likely because in some cases the longer length of Nanopore reads will provide better specificity, however in other cases these longer reads may also contain errors which may lessen specificity, thus this depends on the sequence distance between two closely related species. Comparing MinION and Illumina species abundance resulted in a Pearson correlation coefficient of 0.91 (**Fig. 1b**), which is comparable to previously published work comparing mock community composition using Illumina technology^33^.

**Table 1:** Nanopore flow cell versions and yields. Yield and length metrics for Nanopore runs. Flow cell for sample P8 (2D) was used for two experiments: (a) Initial Metrichor basecalled run which was abandoned due to network lag and (b) Local basecalled run which was used for subsequent analysiss. Runs 1-8 were performed using an earlier version of MinKNOW (ONT’s control software) which classified reads into “pass” and “fail” according to quality. Runs 9-14 used a later version of MinKNOW which removes the quality distinction, however some reads can still fail basecalling. The low yield for run 10 (P49A) is due to the flow cell having a low number of active pores at QC.

| Run | Sample | Flow cell | Seq'ng kit | Library type | Total no. of reads | No. of pass reads | Mean length of pass reads (bp) | Pass read N50 (bp) | Longest pass read (bp) |
| --- | --- | --- | --- | --- | --- | --- | --- | --- | --- |
| 1 | Mock | R7.3 (MAP103) | MAP006 | 2D | 148,441 | 71,675 | 3,047 | 5,497 | 40,561 |
| 2 | P10N | R7.3 (MAP103) | MAP006 | 2D | 145,342 | 82,734 | 2,926 | 3,910 | 17,979 |
| 3 | P10R (1) | R7.3 (MAP103) | MAP006 | 2D | 103,705 | 35,560 | 2,958 | 3,967 | 18,069 |
| 4 | P10R (2) | R7.3 (MAP103) | MAP006 | 2D | 45,486 | 12,118 | 3,832 | 4,832 | 23,423 |
| 5 | P10V (1) | R7.3 (MAP103) | MAP006 | 2D | 165,026 | 53,437 | 925 | 1,087 | 11,300 |
| 6 | P10V (2) | R7.3 (MAP103) | MAP006 | 2D | 69,427 | 21,780 | 2,027 | 2,318 | 15,206 |
| 7 | P8 | R9.4 (MIN106) | LSK208 | 2D | (a) 787,601<br>(b) 423,422 | (a) 407,818<br>(b) 230,494 | (a) 606<br>(b) 622 | (a) 1,768<br>(b) 1,052 | (a) 18,769<br>(b) 18,095 |
| 8 | P8 | R9.4 (MIN106) | RAD002 | 1D Rapid | 69,442 | 44 | 1,480 | 3,089 | 7,848 |
| 9 | P8 | R9.5 (MIN107) | LSK108 | 1D Ligation | 1,369,544 | basecalling stopped at 833,236 | 1,838 | 3,479 | 897,734 |
| 10 | P49A | R9.5 (MIN107) | LSK208 | 1D Ligation | 84,527 | 72,814 | 1,046 | 1,338 | 34,975 |
| 11 | P205G | R9.5 (MIN107) | LSK208 | 1D Ligation | 2,745,619 | 2,415,324 | 966 | 1,102 | 11,619 |
| 12 | P106I | R9.5 (MIN107) | LSK208 | 1D Ligation | 593,183 | 443,299 | 536 | 533 | 3,655 |
| 13 | P116I | R9.5 (MIN107) | LSK208 | 1D Ligation | 422,155 | 392,783 | 464 | 466 | 20,356 |
| 14 | P103M | R9.5 (MIN107) | RAD004 | 1D Rapid | 1,355,250 | 1,221,699 | 1,484 | 1,957 | 153,729 |

**Figure 1.**
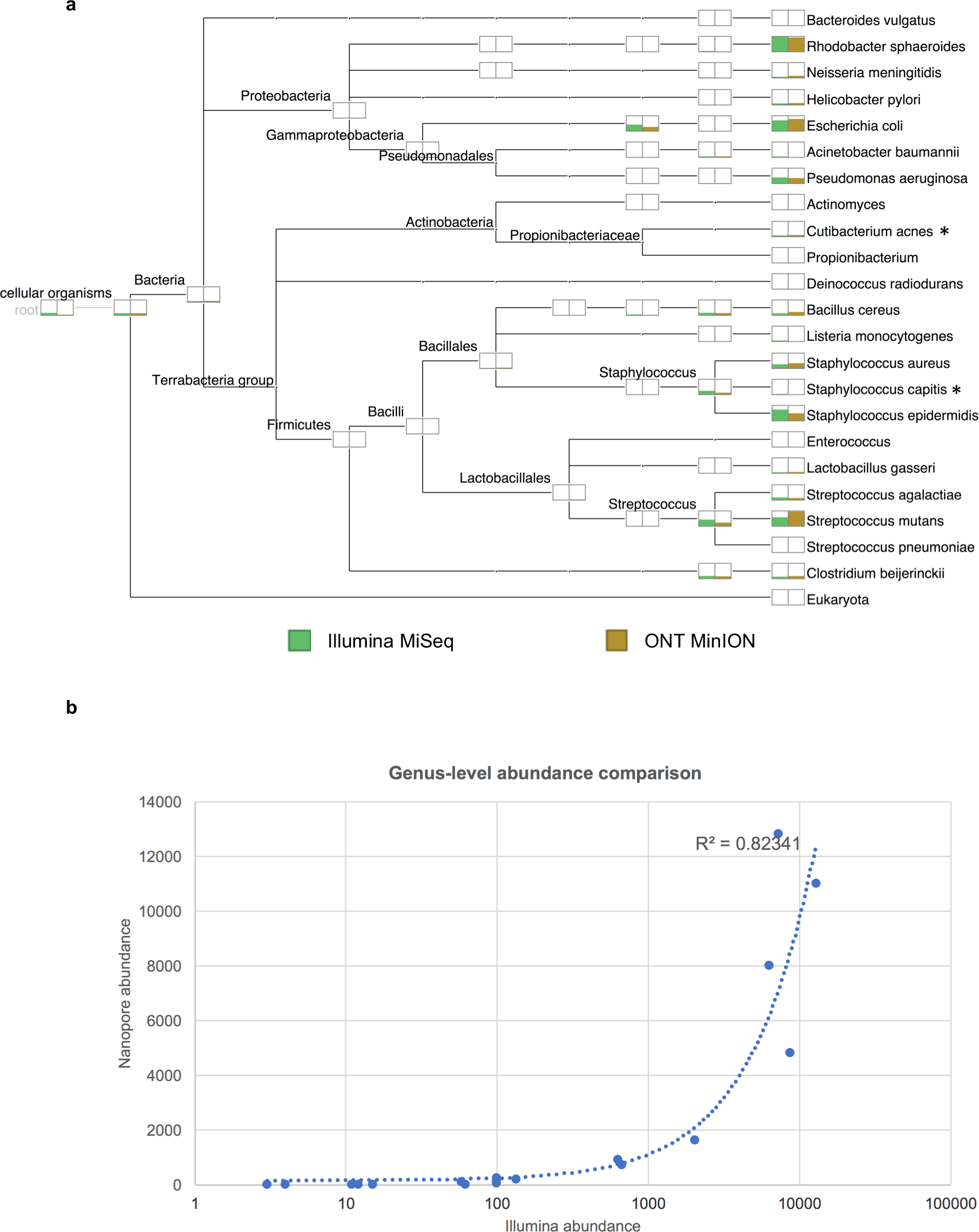
Sequencing of microbial mock community (HM-277D, BEI Resources) using Illumina MiSeq and Oxford Nanopore Technologies MinION sequencing. (**a**) MEGAN taxonomy tree representing species assigned from the mock community control sequenced by Illumina MiSeq (green) and ONT MinION (brown). The height of the bars represents the number of reads assigned for each species taxa. (s*) represents species assigned by Megan but not specified as members in the microbial mock community. (**b**) Genus level abundance correlation plot, with log-based x-axis. Points represent abundance in Nanopore vs. Illumina. Calculated Pearson coefficient 0.91

### Monitoring microbial disturbances in the preterm gut microbiota using MinION

We next sought to determine the relevance of MinION technology in a clinical context; profiling at-risk preterm infants. We performed sequencing on seven infants (three healthy, and four diagnosed with suspected sepsis or NEC, **Supplementary Fig. 1-2, Fig 3a**), blindly selected from a larger ongoing clinical study (BAMBI-BMC genomics^34^). Analysis of faecal samples using PCoA indicated three distinct clusters, which appear to be driven by the presence of either *Bifidobacterium*, *Enterobacter* or *Klebsiella* (**Fig. 2**). We also studied longitudinal samples from one preterm patient (P10) at days 13, 28 and 64 after birth (samples P10N, P10R and P10V respectively, **Fig. 3a**), and compared MinION (version R7.3) to Illumina shotgun and Illumina 16S rRNA gene amplicon sequencing. We initially ran one flow cell for P10N, but it was necessary to run two flow cells for P10R and P10V to generate sufficient yield on this earlier MinION chemistry. This study generated between 145,342 and 234,453 reads per sample across 5 flow cells (**Table 1**). We confirmed that the MinION sequencing depth was sufficient to capture the complete species diversity of the samples, by computing rarefaction curves (**Supplementary Fig. 3**). For all sequencing technologies and samples, the vast majority of species diversity was captured by ~20,000 reads. This analysis indicated that the yield and accuracy of R7.3 MinION flow cells was sufficient for analysis of low bacterial diversity that is characteristic of preterm infant samples^35^.

**Figure 2.**
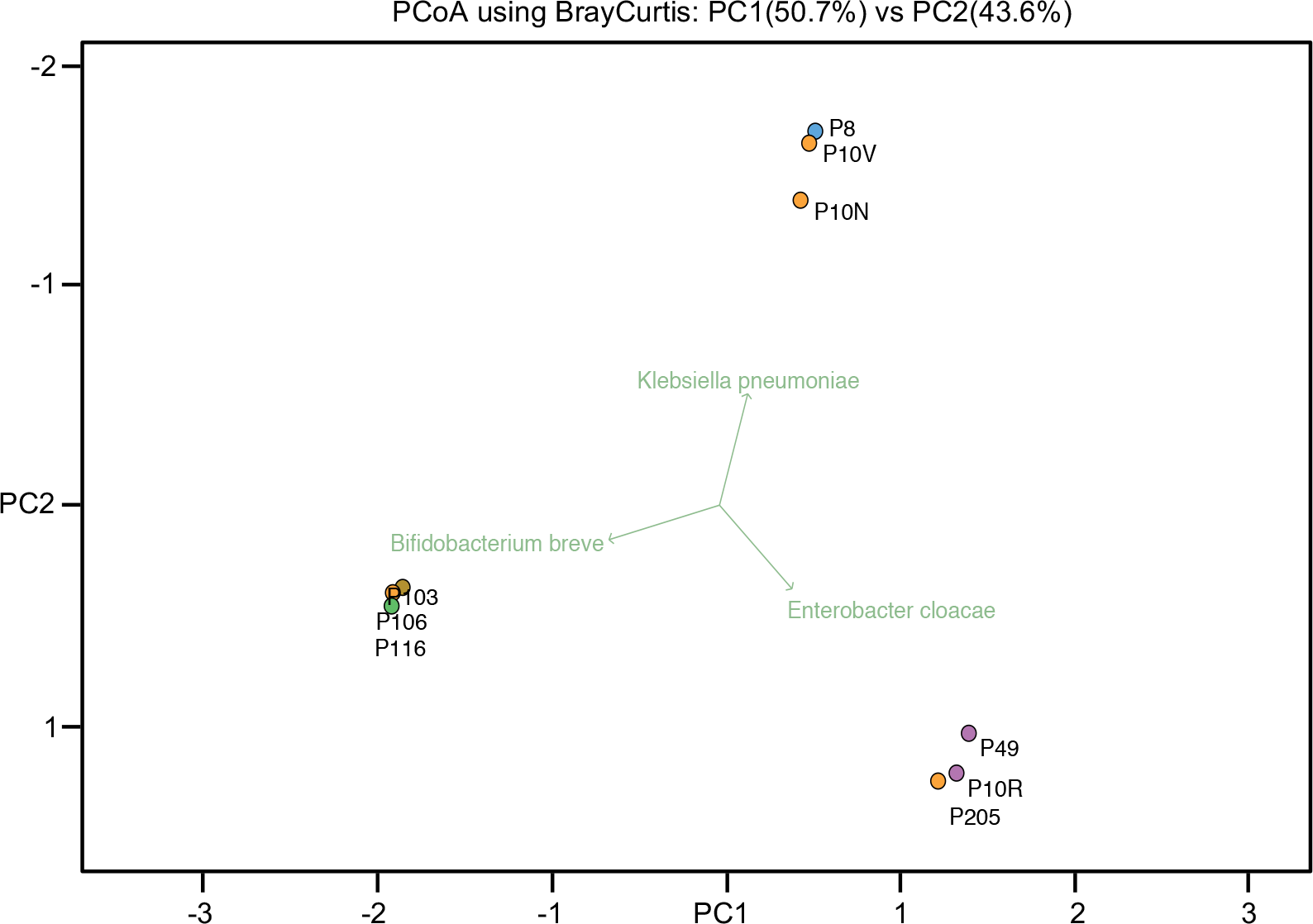
Principal Coordinate Analysis (PCoA) based on taxonomic assignments obtained using MinION nanopore on preterm faecal samples. PCoA was performed based on the bacterial community profiles from profiling faeces from preterm infants. Samples used for this plot were classified into two groups: (i) healthy preterms (P106, P103 and P116) and (ii) preterms diagnosed with NEC or sepsis (P8, P49, P205 and P10). PCoA plot indicates that distribution of samples from healthy preterms was distinct from samples belonging to infants diagnosed with suspected sepsis or NEC.

**Figure 3.**
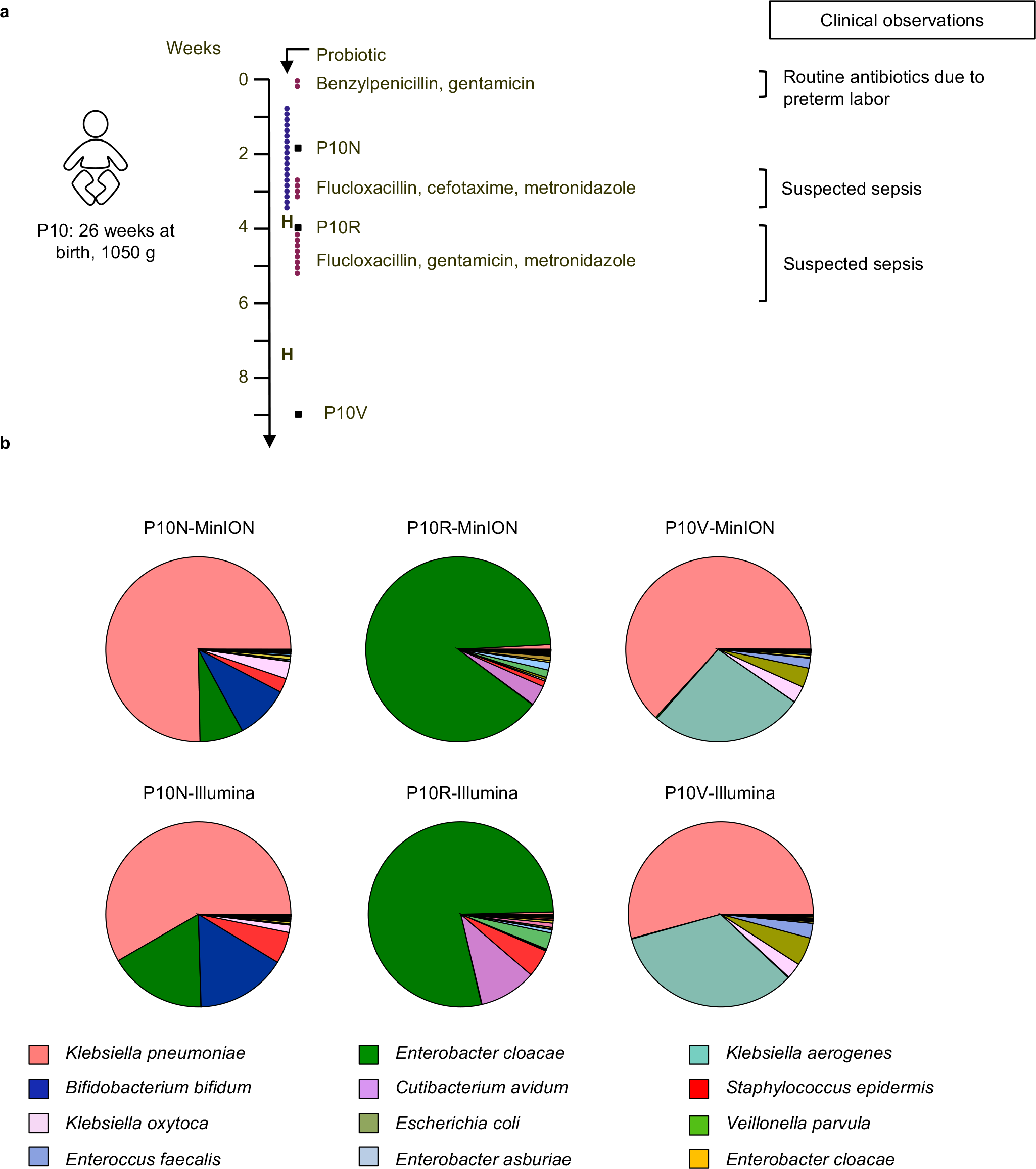
Longitudinal study on preterm P10 using MinION and Illumina sequencing. (**a**) Timeline diagram of preterm P10 indicating times of faecal sample collection (P10N, P10R and P10V), duration of antibiotic and probiotic treatment, and relevant clinical observations. The timeline diagram is divided into weeks and dots represent days within the scale. Blue dots represent days of probiotic treatment, red dots antibiotic treatment, black squares time points for sample collection and letter H transfer of the preterm to another hospital. (**b**) Sequencing data from ONT MinION and Illumina HiSeq 2500 from preterm P10. Pie charts represents taxonomic profiles at different time points P10N, P10R and P10V assigned by MEGAN. The top row corresponds to results obtained using MinION sequencing and bottom row displays results using Illumina HiSeq. Figure legend comprise the 12 most abundant taxonomic taxa obtained. Further information on all the bacteria taxa and the number of reads obtained can be found in **Supplementary Table 3**.

When we compared MEGAN’s taxonomic assignments obtained using MinION vs. Illumina data we again observed that the results were comparable for the majority of the bacterial genera present in preterm infant P10 (e.g. *Klebsiella*, *Enterobacter*, *Veillonella*, *Staphylococcus* and *Bifidobacterium*) (**Fig. 3b**). Interestingly, for sample P10N (collected when the infant was receiving probiotic supplementation) both platforms indicated the potential presence of one probiotic strain, *Bifidobacterium bifidum*. MinION and Illumina also detected the presence of *Enterobacter cloacae* (in sample P10R), a notorious pathogen causing late onset sepsis (LOS) in preterm infants^36^, which correlated with the clinical diagnosis of suspected sepsis in this patient (**Fig. 3a**, however no clinical microbiology testing was carried out to confirm this). Corresponding 16S rRNA gene data (**Supplementary Fig. 4**) provided similar profiles to our shotgun results (albeit with different abundances), but unsurprisingly the short 16S rRNA gene reads failed to differentiate some bacteria taxa even at genus level e.g. members of the family *Enterobacteriaceae*, which comprises commensal gut bacteria as well as opportunistic pathogens and whose full-length 16S rRNA genes are often indistinguishable from one another. Overall, these data highlight the potential for MinION technology and the shotgun sequencing approach in a clinical setting to confirm the impact of microbiota intervention studies (i.e. probiotics) and elucidate potential causative pathogens which may lead to clinical infection diagnoses.

Administration of antibiotics can lead to disruption of early colonisation by gut microbes and create a selection pressure that may also change the profile of AMR genes; the ‘resistome’^37^. We also determined the AMR profile in preterm infant P10 using the Comprehensive Antibiotic Resistance Database (CARD) and compared MinION sequencing data to HiSeq Illumina data. **Supplementary Fig. 5** represents a summary of the AMR genes detected using both sequencing technologies. Overall, if we classify the AMR genes according to mode of action, the detection efficiency of MinION and Illumina was comparable, with only four genes with unique resistance mechanisms (*mphC*, *fusB, sat-4 and vanRG*) detected exclusively by Illumina. Focusing on gene abundance, four main groups: efflux pumps, β-lactamases, aminoglycosides and fluoroquinolones were particularly prevalent (**Supplementary Fig. 6**). Importantly, MinION technology was able to detect AMR genes specific for certain species such as *ileS* encoding for mupirocin^38^ resistance in *Bifidobacterium* (sample P10N), or *fosA2*^39^ encoding for fosfomycin resistance, which is specific for *E. cloacae* (sample P10R, **Supplementary Table 2**). These data illustrate the ability of MinION technology to detect the pool of AMR genes present in the samples tested, and determine species-specific AMR genes.

Infant P10 samples were initially profiled using the older flow cells (R7.3), which provided comparable data to Illumina, we subsequently ran additional preterm samples using more recent MinION flow cells (R9.4), to measure the effect of significant improvements in yield, accuracy and reproducibility. Taxonomic profiling of healthy infants (P106 and P116) indicated a dominance of *Bifidobacterium (B. breve and B. bifidum,* **Supplementary Fig. 7a and 7b**), with presence of *B. bifidum* correlating with the supplemented probiotic species. AMR profiling of these samples using MinION sequencing data revealed a limited ‘resistome’, consistent with the taxonomic profiles and the short antibiotic treatment given (**Supplementary Fig. 7c and 7d)**. Mupirocin resistance was detected, which is a common resistance profile in *Bifidobacterium*. Interestingly levels of other common multidrug resistance determinants were very low in these samples (e.g. ß-lactamases), which may correlate with the low abundance of potentially pathogenic bacteria. These data indicate that recent advances in MinION technology can also successfully profile preterm gut metagenomic samples, including determination of a known species (i.e. *B. bifidum*), and AMR profiles.

### New bioinformatics tools utilise MinION specific features for improved rapid characterisation of gut-associated pathogenic bacteria and resistance profiles

As described above, MinIONs provide near real time sequencing and longer reads than Illumina, but new software is needed to take advantage of these powerful features. ONT’s subsidiary Metrichor provide a cloud-based classification tool called What’s In My Pot? (WIMP). We initially trialled this, but found analysis lagged behind sequencing and the lack of user control over database and classification tool was restrictive for our application. Additionally, the service is now operated commercially. To improve speed and incorporate bespoke analyses, we added real-time functionality to the NanoOK software^30^ creating the new NanoOK RT tool. NanoOK RT takes the sequence reads as they are generated by the ONT software, batches them up for efficiency, and then aligns them to bacterial and CARD databases. A second tool, NanoOK Reporter, provides a graphical user interface which allows the user to view community composition (as a list, donut plot or taxonomic tree) assigned by the Lowest Common Ancestor approach, reports the most prevalent AMR genes, and takes advantage of long reads to perform ‘walkout’ analysis from the AMR gene into flanking DNA of the host bacteria containing the genes.

Having successfully detected a specific beneficial bacterium (i.e. *B. bifidum* after probiotic supplementation) and a potential opportunistic pathogen (i.e. *E. cloacae*) using MinION technology, we next profiled samples from three infants that were clinically diagnosed with NEC at the time point of sample collection (**Supplementary Fig. 2**). Samples were sequenced using R9.4 flow cells and data analysed using NanoOK RT. Infants P49 and P205 samples both contained high proportions of *E. cloacae* (**Fig. 4a** **and 4c**), Analysis of the ‘resistome’ of these infants highlighted a significant number of AMR genes and classes, particularly in P205 (i.e. aminoglycoside resistance and ß-lactamases), which were detected within minutes of sequencing start (**Fig. 4b** **and 4d**). Although these babies had *E. cloacae* dominated gut microbiota profiles, and a significant community ‘resistome’, they also harboured other potentially pathogenic bacteria, highlighting the clinical importance of determining which bacteria are harbouring AMR genes for downstream treatment options.

**Figure 4.**
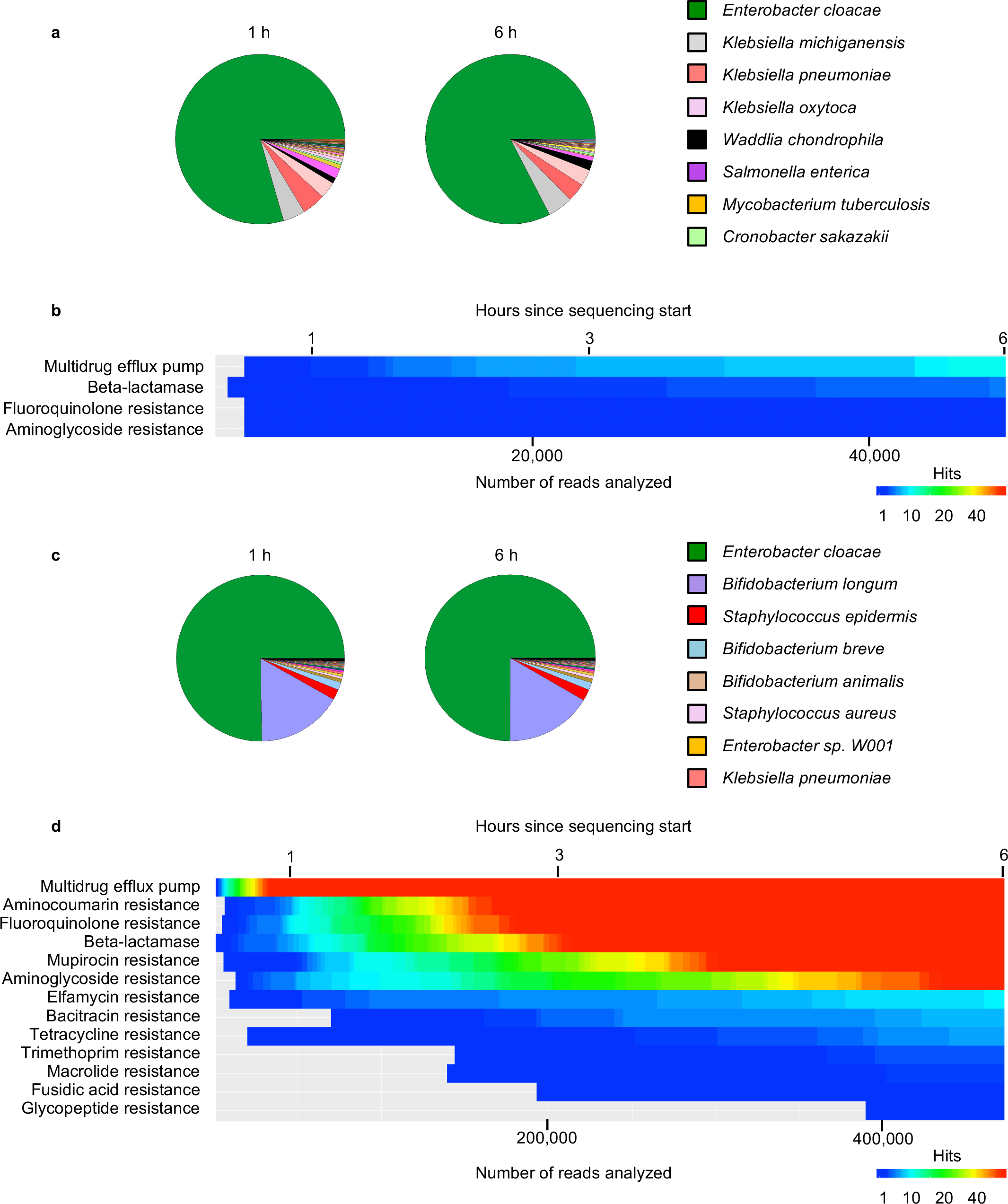
Rapid diagnostic using MinION technology for preterms clinically diagnosed with suspected NEC (P49 and P205). (**a**), (**c**) Taxonomic profiles comparing results obtained at 1h and 6h since sequencing started. Results for preterm P49 are highlighted in **a** and for preterm P205 are highlighted in **c**. Figure legends comprise the 8 most abundant taxonomic taxa obtained. Further information on all the bacteria taxa and the number of reads obtained can be found in **Supplementary Table 3**. (**b**),(**d**) Heat maps displaying number of CARD database hits detected among the most common groups of antibiotic resistance genes found in preterm P49 and P205, respectively. Top and lower panel indicate the hours since sequencing started and the number of reads analyzed. Further information on all the AMR genes obtained can be found in **Supplementary Table 5**.

As MinION reads are typically longer than Illumina reads we reasoned we could extract additional information by examining flanking sequences either side of each AMR hit and searching the NCBI nt database for hits that were independent (defined as ≥ 50 bp) from the AMR sequence. Using this ‘walkout’ approach, which is available through a button click in the NanoOK RT tool, we determined that for infant P205 the vast majority of AMR genes mapped back to *E. cloacae* (88%, **Fig. 5a**), which was also the dominant species taxonomically, with the remaining resistance genes (6%) associated with *B. longum* (i.e. mupirocin resistance). Contrastingly, although infant P49 had very similar levels of *E. cloacae* compared to infant P205, only 60% of AMR hits were associated with *E. cloacae, and* although *Klebsiella* represented a small proportion taxonomically in P49, *Klebsiella* species (*K. pneumoniae*, *K. michiganensis* and *K. oxytoca*) appeared to encode for a range of AMR genes (e.g. OXA-2 (ß-lactamases), CRP and mexB (efflux pumps) and patA and mfd (fluoroquinolone resistance)), making up >30% of total AMR genes present in this infant sample (**Fig. 5** and **Supplementary Table 5**). These data, highlight that MinION sequencing data coupled with the NanoOK Reporter analysis software is able to map specific AMR determinants to specific pathogenic bacteria, and could facilitate more appropriate treatment.

**Figure 5.**
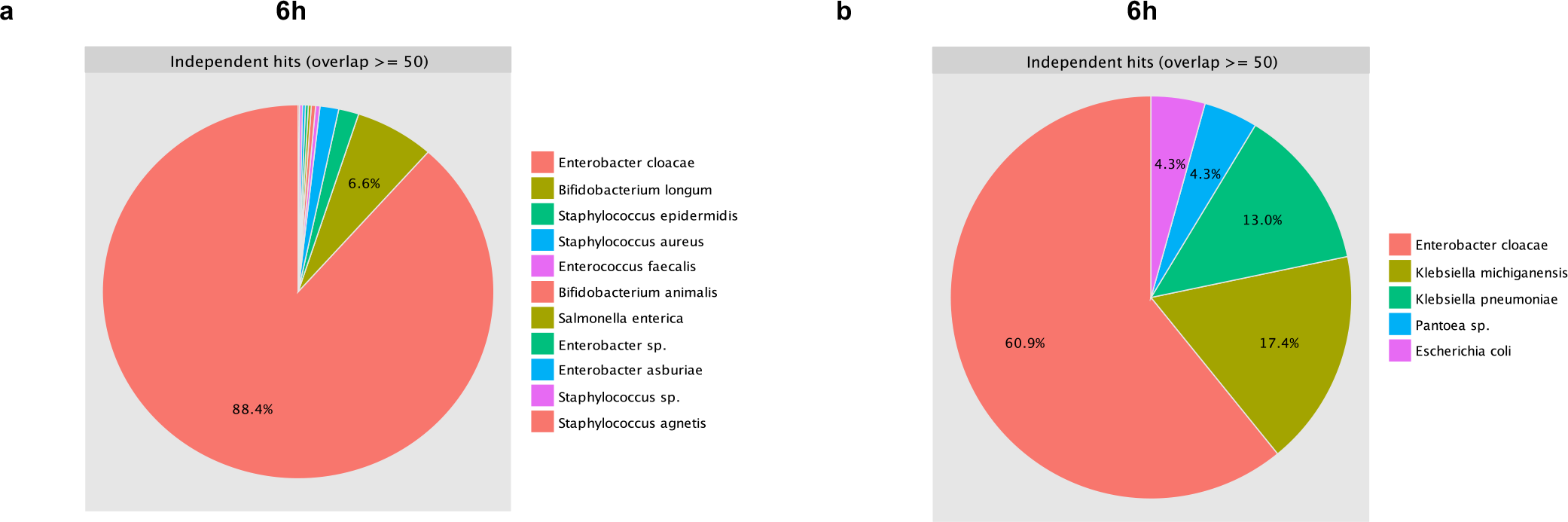
Walkout study for preterms P205 and P49 reported by NanoOK RT software. (**a**) Results from independent hits (defined as ⋝ 50 bp overlap from the AMR sequence) at 6 hours of sequencing for preterm P205. (**b**) Results from independent hits (defined as ⋝ 50 bp overlap from the AMR sequence) at 6 hours of sequencing for preterm P49.

Next we sought to perform a ‘real time’ run to evaluate how rapidly the MinION coupled to the new NanoOK RT software could provide relevant data to help pathogen diagnosis and guide antibiotic treatment. Current rapid clinical microbiology tests, including determining antibiotic susceptibility, take between 36 and 48h. We performed a ‘real-time’ run (from sample preparation to data analysis) using MinION R9.4 flow cells on a faecal sample from a preterm infant (P8) clinically diagnosed with suspected NEC (**Supplementary Fig. 2c**). Notably, this infant was exposed, before sample collection, to 43 days of non-concurrent antibiotic treatments (i.e. benzylpenicillin, gentamicin, meropenem, tazocin, vancomycin, flucloxacillin, metronidazole and amoxicillin). We initially performed this run timing all stages used for this pipeline including sample preparation (90 min), DNA quality control (45 min), 2D MinION library preparation and loading onto the MinION flow cell (2 h), and sequencing-and-data analysis (40 min for first non-specific AMR hit, 4 h 55 total), although we did have issues with basecalling which necessitated restarting. Taxonomic results, AMR genes detected and walk out results for this run can we seen in our initial preprint^40^.

Subsequently, we performed another ‘real-time’ run on the same sample, this time utilising 1D rather than 2D libraries (2D is now phased out by ONT), and generated 1.37 million reads in a full 48 hour run. The removal of an incubation step in a simplified library protocol resulted in saving approximately 15 minutes compared to the 2D library, thus it took approximately 4 hours to go from sample receipt to sequencing (**Fig. 6**). The sequence reads were analysed using the NanoOK RT pipeline (see Online Methods for further details). The first 500 reads immediately indicated a dominance (332 reads) of *K. pneumoniae* (a potential causative organism that has been associated with NEC pathogenesis in preterm infants^41^), as well as *Proteus mirabilis* (30 reads). By 1 h after sequencing start (5 h total), the pipeline had analysed 20,000 reads and *K. pneumoniae* accounted for around 70% of reads. These reads were much longer (N50 3,479 bp) than the previous 2D run (N50 1,052 bp), meaning that at the 1h point NanoOK RT had analysed over 3× more sequence data in this new 1D run. To further verify we had sequenced enough of the bacterial diversity existing in the sample at this time point, we compared taxonomic profiles from analysis completed at 6 h, time-point chosen due to clinical relevance to NEC deterioration (101,000 reads, 10 h total time). This comparison verified that there were no significant qualitative differences between the two taxonomic profiles (**Fig. 7a**).

**Figure 6.**
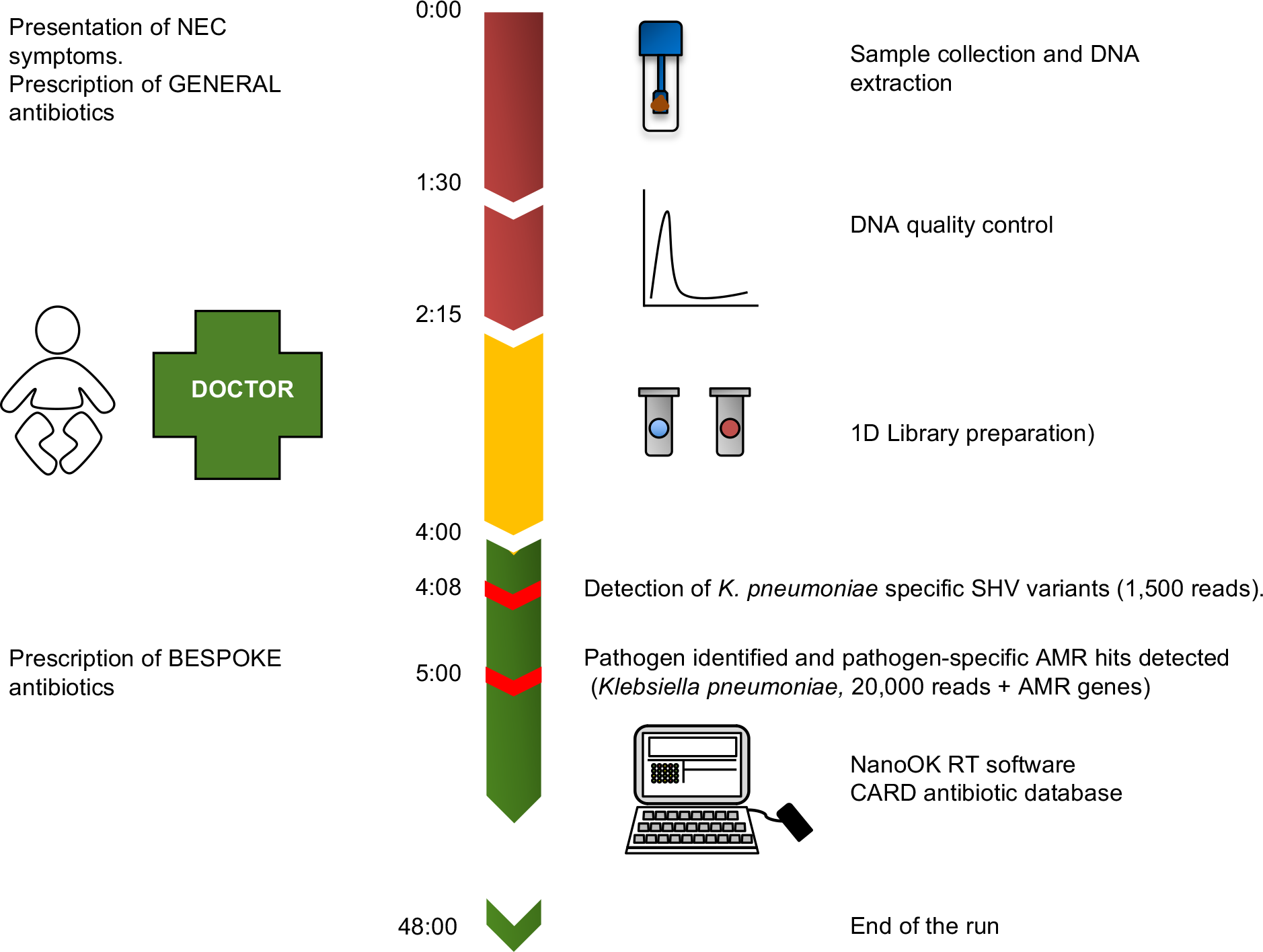
Timeframe diagram for ‘real time’ run performed for rapid diagnostic of a preterm infant (P8) suffering from NEC. Step 1 (red, 2h 25min): Sample collection, DNA extraction and quality control. Step 2 (yellow, 1h 45min): 1D library preparation incorporating bead clean up and DNA repair. Step 3 (green): data analysis using local base calling and NanoOK RT. Pathogen detection (*Klebsiella pneumoniae*) and *K. pneumoniae* specific AMR genes were first detected at 4 hours and 8 minutes (1,500 reads analysed). Left side of the panel indicates clinical symptoms and general guidelines for antibiotic prescription.

**Figure 7.**
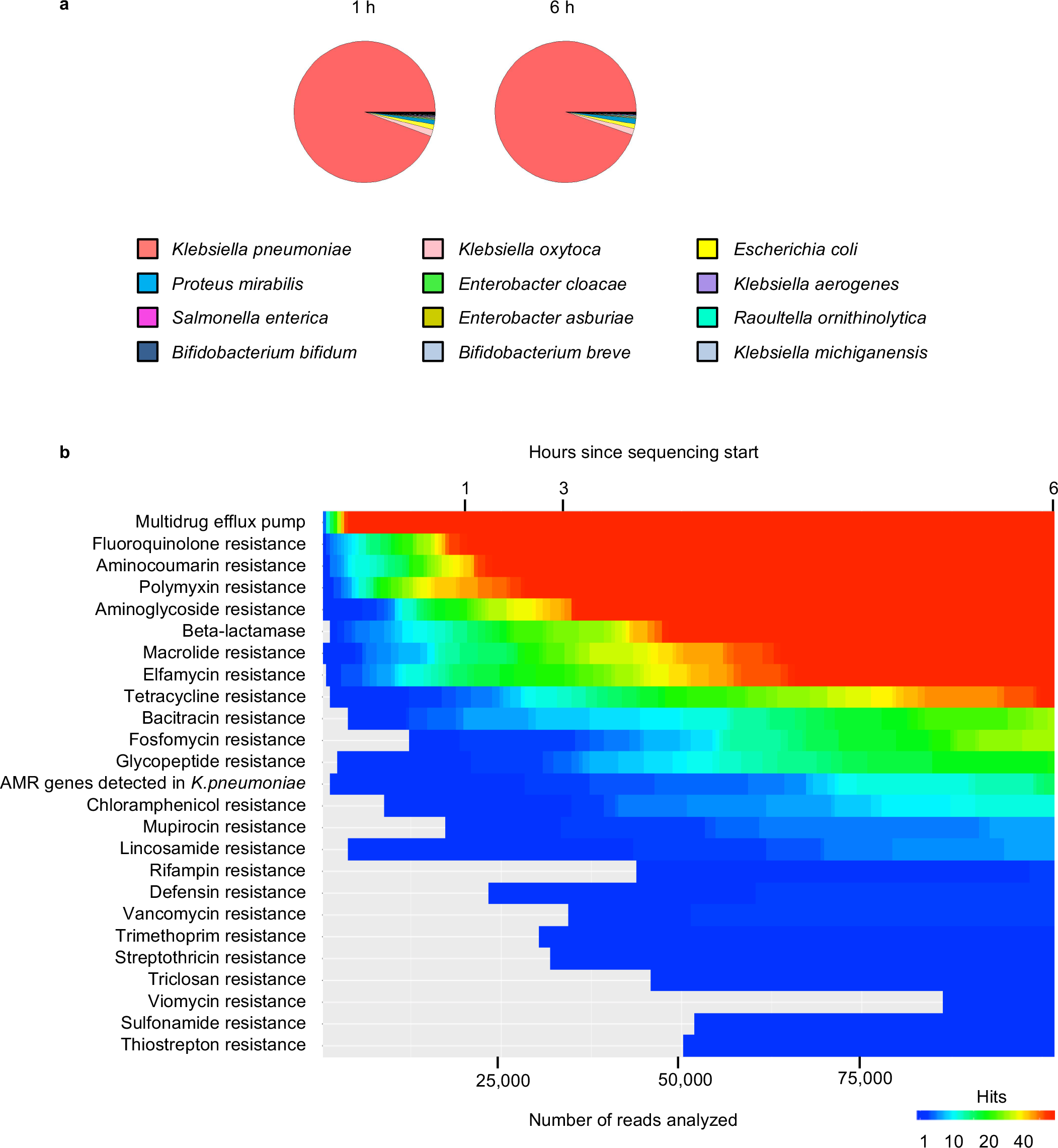
Rapid diagnostic of preterm P8 clinically diagnosed with suspected NEC. (**a**)Taxonomic profiles obtained using MinION technology at 1h, and 6h since sequencing started. Figure legend comprises the 12 most abundant taxonomic taxa obtained. Further information on all the bacteria taxa and the number of reads obtained can be found in **Supplementary Table 3**. (**b**)Heat map displaying number of CARD database hits detected among the most common groups of antibiotic resistance genes found in preterm P8. Top and lower panel indicate the hours since sequencing started and the number of reads analyzed, respectively within this timeframe. Further information on all the AMR genes obtained can be found in **Supplementary Table 5**.

As highlighted previously, it is clinically important to detect AMR genes in metagenomic samples from preterm infants to guide appropriate antibiotic prescription. In our ‘real-time’ run we determined how rapidly we could map AMR genes to the CARD database. **Fig. 7b** shows the huge number of AMR gene classes detected throughout the run, including polymyxin, aminoglycoside, tetracycline, quinolone resistance, β-lactamases and efflux pumps, all of which were detected in as little as 1 h after sequencing start. We were able to detect *K. pneumoniae*-specific SHV variants^42^ as early as 6 min (at 1,500 reads, 4 h 8 min total time), whilst other lower abundance AMR genes in the sample, such as those conferring trimethoprim, sulphonamide and streptothricin resistance, were not detected until 3-4 h post sequencing (7-8 h total). Interestingly, the 1D, compared to our initial 2D run produced significantly more sequence data, which also allowed detection of a substantially greater number of AMR genes, which could be used in downstream clinical decisions.

We also used NanoOK Reporter to perform AMR ‘walkout’ analysis on this infant sample, which indicated that the majority (~90%) of AMR genes within the whole metagenomic sample mapped to *K. pneumoniae* (**Fig. 8**), including multidrug exporters such as *acrB* or *oqxA*, conferring resistance to tetracycline, chloramphenicol, and fluoroquinolones, *vanSC* (resistance to vancomycin), *tet 41* (resistance to tetracycline) and *dfrA20* (resistance to trimethoprim). We also correlated specific AMR gene cassettes to *P. mirabilis* including OXA-63, which can confer cephalosporin resistance, *and tet34 resistance to tetracycline* (**Supplementary Table 5**).

**Figure 8.**
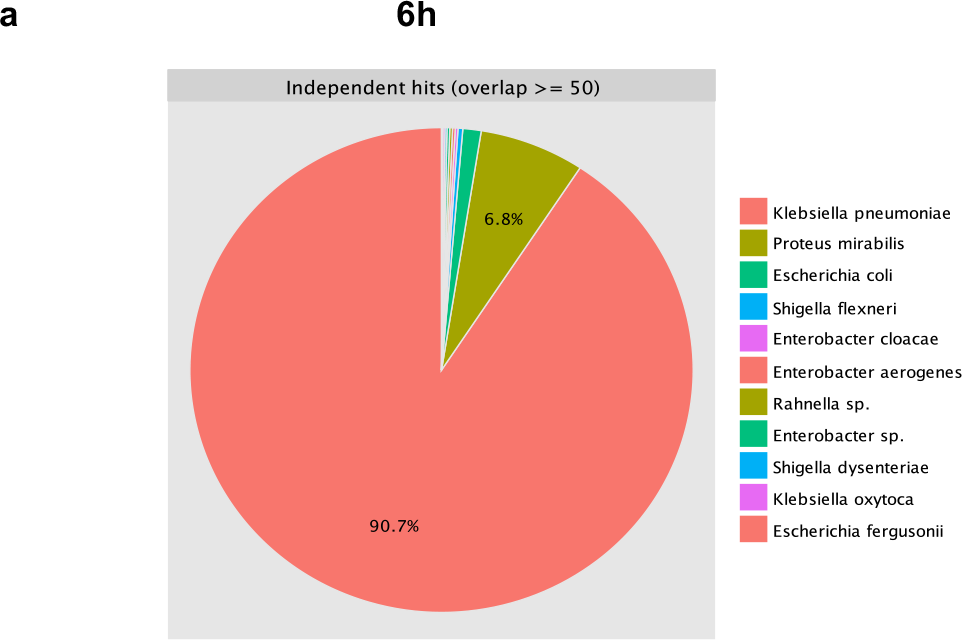
Walkout study of preterm P8 reported by NanoOK RT software. (**a**) Results from independent hits (defined as ⋝ 50 bp overlap from the AMR sequence) at 6 hours of sequencing.

### Phenotypic characterisation of *Klebsiella pneumoniae* isolates from preterm infant P8

To validate the genotypic data obtained from our ‘real time’ MinION nanopore run, we isolated nine *K. pneumoniae* colonies from patient P8, and performed alignment analysis on their 16S rRNA gene sequences which showed similarity levels ranging from 99.8% to 100% (**Supplementary Table 6**). To further confirm our metagenomics findings, we performed Illumina and MinION whole genome shotgun sequencing and assembly on one *K. pneumoniae* isolate. The Illumina assembly resulted in 69 contigs totalling 5.73Mb, with an N50 of 310,421, while the nanopore assembly produced a single contig of 5.47Mb and two further contigs totalling an additional 0.37Mb. AMR genes including, *FosA* (encoding fosfomycin resistance), *acrA, oqxA*, and *oqxB* (encoding efflux pumps), and *SHV-185* (encoding extended-spectrum β-lactamases, ESBLs), correlated between whole genome sequencing data and walk-out analysis from the metagenomic samples (**Supplementary Fig. 8**). Interestingly, analysis with NanoOK RT indicated a wider variety of AMR genes in *K. pneumoniae* conferring resistance towards antibiotics this infant received e.g. *OXA* genes (resistant to benzylpenicillin and amoxicillin), *imiS, NDM-9* and *THIN-B* genes (resistant to meropenem), aminoglycoside acetyltransferases (*aac*) and aminoglycoside-phosphotransferases (*aph*) (resistant to gentamicin) and *vanRI*, *vanTG* and *vanTmL* (resistant to vancomycin). Overall these results indicate MinION metagenomic sequencing using NanoOK Reporter and ‘walkout’ analysis is faster and provides robust clinically relevant AMR data that may help guide antibiotic treatment.

In order to demonstrate antibiotic resistance phenotypes correlating to presence of AMR genotypes we tested the susceptibility of the same *K. pneumoniae* isolate with the seven most commonly used antibiotics in NICUs (**Supplementary Fig. 9**). Interestingly, the *K. pneumoniae* isolate was found to have a higher minimum inhibitory concentration (MIC) breakpoint value for those antibiotics that were prescribed to the preterm infant P8 i.e. benzylpenicillin, amoxicillin, metronidazole, gentamicin and vancomycin. In contrast, the only MIC breakpoint value lower than those put forward by the European Committee on Antimicrobial Susceptibility Testing (EUCAST^43^) was for cefotaxime, an antibiotic not prescribed to the infant at sample collection point. Notably, these data also correlate with the AMR data generated by NanoOK reporter and the ‘walkout analysis’, with the only exception of metronidazole resistance which was only detected after whole genome sequencing of the isolate (gene *msba*, **Supplementary Fig. 8**).

### Further enhancements using rapid sequencing technology

ONT now produce a rapid library kit which takes as little as 10 minutes preparation time. We had previously tried an older version of this kit (SQK-RAD002) but had not been able to get good quality data from infant stool samples. Recently, a new version (SQK-RAD004) was released which we evaluated using a sample from healthy infant P103. Profiling the gut microbiota of infant P103 produced 1.2 million reads (read N50 of 1,957 bp, longest read mapping to nt database with length of 17,069 bp), with a patient sample to analysis time around 60 minutes faster than our 1D ‘real time’ run on infant P8. This was a healthy infant, with no infection diagnoses, and we confirmed a dominance of commensal *Bifidobacterium* species, including *B. bifidum* (also probiotic species, **Supplementary Fig. 10a**). AMR profiling and walkout analysis using NanoOK RT, followed the same trend as healthy infants P106 and P116 previously analysed in this study, and indicated a high proportion of mupirocin resistance (specific to *Bifidobacterium*) and tetracycline resistance (**Supplementary Fig. 10b)**. Utilisation of this new methodology enhanced the speed and overall data generation for preterm microbiota samples, thus further indicating the utility of using MinION for these types of clinical samples.

Due to the amount and quality of the sequence data generated during this run, we also performed a reference-guided assembly of the *B. bifidum* genome from this metagenomic data which resulted in 3 contigs with an average identity of 98.86%. (**Supplementary Fig. 11)**. This clearly demonstrates the potential of nanopore sequencing to resolve whole genomes out of metagenomic samples, though the error rate is still currently high enough to make SNP analysis challenging.

## DISCUSSION

With widespread concerns about increasing AMR rates worldwide, there is a pressing clinical need for tailored (rather than broad spectrum) antibiotic use. Achieving this goal requires faster more detailed microbial diagnostics which should also lead to better patient outcomes. Gold standard microbial laboratory screening is limited by the growth rates of possible pathogens on selective media and the combinatorial tests that can be conducted. While microbiological tests take days, newer sequencing platforms, e.g. Illumina, could manage this in a faster manner and provide rich data that can also be used to track hospital epidemics^44^, but requires specialised laboratories and large capital investment. The new MinION sequencer seems an ideal tool because it is smaller, faster and cheaper than Illumina machines. Using a combination of improved Nanopore sequencing chemistries, and developing our own open source Nanopore analysis packages, NanoOK RT and NanoOK Reporter, we showed that the MinION platform is able to successfully profile known metagenomes (i.e. mock community), and even clinical samples. Importantly, MinION sequencing data using the new R9.4 flow cells were comparable in discriminatory power to the conventional Illumina sequencing platform and provided clinically relevant information within just 5 h from sample receipt. We were able to demonstrate three clinically relevant, and actionable, pieces of data: 1) presence and abundance of microbial species in the sample, 2) overall antibiotic resistance gene profile and 3) species-specific antibiotic resistance profiles.

Our first findings, using a known metagenomic sample (mock community), indicated that the MinION is suitable for detection of microbes, which is in agreement with a previous study also using R7.3 flow cells^29^, and suggested profiling of mixed clinical samples may be possible. Thus, we next tested longitudinal samples from a preterm infant residing in NICU where improved diagnostic tools could be used to improve outcomes. We found MinION and Illumina analyses were comparable in discriminating species and indicated one of the probiotic strains (i.e. *B. bifidum*) during the supplementation period was present (**Fig. 3b**). Furthermore, these data provided information on probiotic therapy indications after antibiotic administration; absence of these beneficial species may correlate with a ‘disturbed’ microbial profile^45^. Specifically, *E. cloacae* completely dominated the microbiota (sample P10R) and is a known sepsis pathogen (able to disseminate to the blood) in immunocompromised, including preterm patients^46^. Further analysis of other preterm samples from infants P106 and P116 (using R9.4 flow cells) also indicated that a dominant bifidobacterial gut microbiota profile correlated with health, as these infants did not have any clinical diagnosis at sample collection. Furthermore, whilst these samples indicated presence of AMR genes, their ‘resistome’ was markedly reduced in comparison to infants harbouring potentially pathogenic bacteria e.g. *Klebsiella* and *Enterobacter*. Therefore, profiling beneficial microbiota members and opportunistic gut associated pathogens may provide key clinical data to prevent subsequent systemic infections.

Clinical microbiology labs carry out critical tests for the detection of pathogens and their antibiotic sensitivities. This is particularly pertinent due to the increasing prevalence of antibiotic-resistant bacteria and lack of novel antibiotic development. We believe MinION sequencing may provide a rapid tool for crafting bespoke antibiotic treatment strategies in time-critical patients. When we investigated AMR genes detected by MinION and Illumina, both sequencing platforms generated reads mapping to genes with similar antibiotic resistance mechanisms (**Supplementary Fig. 5**), and only 4 genes (*mphC*, *fusB*, *sat-4 and vanRG*) with unique resistance mechanisms out of all 146 AMR genes were detected exclusively by Illumina. This result may be correlated to the lower MinION read count, and so could be mitigated by ongoing improvements in MinION technology. Notably, we observed the presence of AMR genes that corresponded to prescribed antibiotics; β-lactamase and aminoglycoside genes (conferring resistance to benzylpenicillin and gentamicin respectively), while fluoroquinolone resistance genes did not correlate to any prescribed antibiotics, and so may relate to AMR gene transfer of strains from other sources. Thus, AMR profiling may guide clinical treatment decisions at an earlier stage of patient care.

Whilst our comparisons of preterm gut-associated microbial profiles highlighted the significant scope for MinION in a relevant clinical context, we wanted to demonstrate that the entire pipeline of sample preparation, library construction, sequencing and analysis could be carried out rapidly and in ‘real-time’. Importantly, we used the most recent flow cells (R9.4) for this study, which have an improved error rate (~5-7% for 1D) and yield. Specifically, for this study we added significant functionality to NanoOK for the ‘real-time’ analysis of species abundance and antibiotic resistance genes. We initially examined two ill infants, P49 and P205, both of which had high levels of *E. cloacae*, and a significant resistome, which may correlate with their clinical diagnosis of suspected NEC, similar to infant P10, who was diagnosed with suspected sepsis. Importantly, additional analysis using the new NanoOK Reporter functionality, enabled us to determine what specific taxa harboured these AMR genes, and efflux pumps including *E. cloacae*, which was the dominant species present. Furthermore, although *Klebsiella spp.* represented a more minor component of the P49 microbial community they appeared to harbour > 30% of the total AMR genes present. Thus, in patient P49 a poorly chosen antibiotic treatment could target *Klebsiella* and miss the predominant *E. cloacae* or could target only *E. cloacae* lead to an increase in *Klebsiella* pathogenic species, whereas the best treatment would target both sets of pathogenic species. Performing a walkout, rather than *de novo* metagenomic assembly is considerably less compute intensive, and critically is far faster and more accurate, a very important consideration in the clinical environment. These data indicated that relevant AMR genes detected in a metagenomic sample and further mapped to known pathogenic species may facilitate tailored antibiotic treatment strategies for critically ill patients. Additionally, using MinION metagenomic data we were able to assemble *B. bifidum* (P103) using a reference-guided approach as this infant had received a known species present in the probiotic supplement.

These studies highlighted the ability of MinION and NanoOK to provide important pathogen and AMR information, thus we sought to mimic a more clinically relevant diagnostic approach by performing a real-time run. The samples for this experiment was from an extremely ill preterm infant P8 (born after 26 weeks’ gestation with a birthweight of only 508 g), who had received multiple courses of antibiotics since birth (46 days of antibiotic treatment out of 63 days of life at sample collection) and presented with clinical NEC observations at the time of sample collection. Whilst MinION allows generation of long reads, DNA extraction to maximise this type of data is more time-consuming, thus in this study we utilised a rapid DNA extraction protocol (including a bead-beating step). Furthermore, previous studies, indicate that incomplete DNA extraction significantly biases metagenomic profiles obtained^34, 47^, which may in turn limit pathogen detection and AMR analysis. Therefore, we acknowledge that it may be possible to improve read length in future (sample P8 had N50 of 3,479 bp) through improved DNA extraction methods.

As highlighted in our previous preprint, initial attempts to real-time sequence infant P8 were unsuccessful, due to problems related to quality of flow cells and base calling. Our third attempt generated impressively high yields (101,000 reads) at only 6 hour of sequencing and 1.37 million reads in a full 48 hour run. Taxonomic profiling in real time revealed a *K. pneumoniae*-dominated profile, after just 1 h of sequencing (~20,000 reads), enabling us to confidently ‘call’ this potential pathogen. This analysis was further strengthened as more sequencing at 6 h, gave almost identical microbial profiles (**Fig. 7a**), as did Illumina sequencing (**Supplementary Fig. 12**). These data therefore suggest that this bacterium may be causative in the clinical diagnosis of this infant i.e. NEC. Importantly, *K. pneumoniae* has been linked to preterm NEC (and is supported by corresponding clinical observations), with overgrowth in the intestine linked to pathological inflammatory cascades, facilitated by a ‘leaky’ epithelial barrier^48^. It should be noted that the single species domination in this sample facilitates early detection at lower read depth and low-level pathogen abundance would require deeper sequencing. Profiling of additional more complex samples from NEC diagnosed infants (i.e. P49 and P205) also indicates distinct and differential microbiota profiles (when compared to P8) also at 1 h post sequencing start (**Fig. 4a** **and 4c**), highlighting how rapid diagnosis of pathogen overgrowth is possible using R9.4 Nanopore flow cells.

Whilst detection of individual pathogens is important, a critical additional requisite is identification of AMR profiles so that tailored antibiotic treatment can be used. Real-time analysis of MinION data using NanoOK RT highlighted the presence of a significant metagenomic ‘resistome’, including presence of colistin resistance, a last resort antibiotic, by the detection of genes *arnA*^49^, *PmrB*^50^ and *PmrC.* We noted the greater the sequencing depth the greater the number of AMR genes detected, although importantly we were able to detect a significant number of AMR genes as rapidly as 1 h after sequencing start, including β-lactamases, quinolone, aminoglycoside, and tetracycline resistance genes (**Fig. 7b).**

*Klebsiella* is of particular concern within an AMR context due to the increasing emergence of multidrug-resistant isolates that cause severe infection, which represent a real threat to patient outcomes^51^. Therefore, to improve the potential for guiding antibiotic treatment decisions we performed the ‘walkout’ study from the AMR gene sequences to determine which bacterial species were carrying which AMR genes. Based on these data, a clinical decision (as determined by neonatologist Dr Paul Clarke, in a blinded fashion) at 1 h after sequencing start (5 h total time) would be continuation of gentamicin and flucloxacillin or cefotaxime treatment (NICE guidance^52^ states antibiotics should be given within 1 h of presentation of signs of sepsis). However, after 6 hours of AMR sequence data (10 h total time), we were able to obtain a more informative overview of the full resistance genes profile (101,000 reads, **Supplementary Fig.8**). At this point our analysis indicated presence of *OXA* genes, *CTX-M* genes and *SHV* genes (encoding resistance to flucloxacillin and cefotaxime), and aminoglycoside-acetyltransferases (*aac*) and aminoglycoside-phosphotransferases (*aph)* (resistance to gentamicin). Presence of extended spectrum β-lactamases (ESBLs) may suggest the infant is moved to meropenem, a carbapenem which is often given as a last resort for neonatal infections^53^. However, genes including *imiS*^54^, NDM-9^55^, or *THIN-B*^56^ (which may encode carbapenemase activity against meropenem) indicates meropenem treatment alone may not be sufficient to clear this *K. pneumoniae* infection, and a combination therapy e.g. meropenem with ertapenem may be required^57^. Overall, information at only 10 hours, which would correlate with presentation of first NEC symptoms, is clinically very relevant and potentially provides important information on the pathogen involved and associated AMR profiles that may guide antibiotic treatment choices.

We sought to further benchmark our MinION pipeline by correlating our metagenomic observations with isolated *K. pneumoniae* strains that had undergone genotypic (WGS) and phenotypic (MIC) analysis. Notably, whole genome shotgun sequencing (both Illumina and MinION) indicated agreement with resistance genes determined by the MinION metagenomic run, but the ‘walkout’ approach indicated a significantly greater AMR profile at only 10 h (**Supplementary Fig. 8**). This may due to the significant number of *K. pneumoniae* isolates in this metagenomic sample and corresponding diverse AMR profiles, which would not be captured in a single isolate WGS analysis (although a previous study has indicated it is possible to multi-plex numerous *Klebsiella* isolates onto a single MinION run^58^). When subjecting strains to MIC testing (current gold standard for profiling AMR), we observed phenotypic resistance to all main groups of antibiotics that had been prescribed to infant P8, including gentamicin (which would have continued to be used at 1h post sequencing AMR hits; **Supplementary Fig. 8**). There was good association between AMR gene sequence detection and MIC testing, i.e. *SHV* and β-lactam antibiotics, *aac* and *aph* genes and gentamicin, and *van* genes resistant to vancomycin which highlights that MinION could be extremely useful for rapid AMR profiling. Only metronidazole resistance was identified via MIC testing, that wasn’t present in the MinION metagenomic analysis, although the corresponding resistance gene was only detected in the isolate whole genome sequencing data (gene *msba*, **Supplementary Fig. 8**). As such we expect that rapid analysis using MinION sequencing, would inform early and more appropriate antibiotic choices for patient care, halting the rapid deterioration observed in critically ill patients. Notably, our final MinION run using the new rapid kit (SQK-RAD004) indicates an even quicker rate to determine actionable clinical data, which may provide the difference between life and death in critical patients including septic or NEC preterm infants. Subsequently, phenotypic testing via standard clinical microbiology labs would further refine clinical management of patients. Importantly, it is clear that preterm infants harbour a significant reservoir of AMR genes within the wider microbiome as well as in potentially pathogenic bacteria such as *K. pneumoniae*, which makes bespoke antibiotic treatment decisions non-trivial. Thus, further highlighting the urgent requirement for comprehensive AMR databases, new antibiotic development and alternative treatments such as novel antimicrobials or microbiota therapies.

We obtained MinION bioinformatics results in under 5 hours. In comparison, standard Illumina MiSeq sequencing (paired 250 bp reads) alone normally takes 39 hours, excluding sample preparation and analysis (https://www.illumina.com/systems/sequencing-platforms/miseq/specifications.html), and PacBio (i.e. long-read sequencing) with the same sample prep and quality as our MinION experiment, using the rapid (4 hour) library method (http://www.pacb.com/wp-content/uploads/2015/09/Guide-Pacific-Biosciences-Template-Preparation-and-Sequencing.pdf), 30 minute diffusion and the shortest sequencing run (30 minute) would take over 7 hours even without base calling or bioinformatics, highlighting the significantly quicker detection potential of MinION in a clinical setting.

Throughout this study we have utilised the latest ONT kits for the MinION, with the recently released RAD kit producing a highly significant number of reads rapidly for infant P103. Notably, a number of products announced by ONT could make Nanopore sequencing in clinical settings even more attractive. The SmidgION is an even more compact device than the MinION that can be connected to a mobile phone. At the other extreme, the GridION X5 and PromethION are respectively capable of running between 5 and 48 flow cells simultaneously, facilitating better economies of scale. Sample preparation is already relatively simple, particularly for 1D libraries, but this is likely to be further simplified by VolTRAX, a compact automated library preparation system from ONT that can manipulate fluids around an array of pixels. ONT have demonstrated 1D library preparation directly from an *Escherichia coli* cell culture entirely with VolTrax (https://nanoporetech.com/publications/voltrax-rapid-programmable-portable-disposable-device-sample-and-library-preparation). Balancing the excitement about new products is that the technology is still relatively immature and is frequently updated, issues that must be addressed for a true diagnostic tool. Even within the time-frame of this study we have observed significant advances in optimisation of MinION technology, and we hope this could lead to a stable platform for large-scale clinical testing.

Whilst we show the results of single patient diagnostics alone, if this platform were to be broadly adopted, even within a single ward or hospital, additional epidemiological analysis should be possible. Transmission chain analysis of hospital-acquired pathogens has been shown using Illumina data^44^, and Nanopore sequence-based epidemiology was used to monitor the West Africa Ebola virus outbreak^59^.

## CONCLUSION

In conclusion, we have demonstrated that MinION technology is a rapid sequencing platform that has the ability to detect gut-associated pathogens that have been associated with potentially life-threatening preterm-associated infections in ‘real time’; identification of specific pathogenic taxa (i.e. *K. pneumoniae and E. cloacae*) and corresponding AMR profiles. Data obtained could allow clinicians to rapidly tailor antibiotic treatment strategies (i.e. change from gentamicin to a combination therapy of meropenem with ertapenem) in a rapid (~6 h decision from sample receipt) and timely manner. The utility of this approach was confirmed when compared to Illumina metagenomic sequencing and isolation and characterisation of *K. pneumoniae* strain including WGS and phenotypic (i.e. MIC) testing. This suggests that MinION may be used in a clinical setting, potentially improving health care strategies and antibiotic stewardship for at-risk preterm infants in the future.

## METHODS

Methods and associated references are available in the online version of the paper.

### Accession codes

The Illumina and MinION read data supporting the conclusions of this article are available in the European Nucleotide Archive (http://www.ebi.ac.uk/ena) under study accession PRJEB22207.

## ACKNOWLEDGEMENTS

RML, DH and MDC’s MinION work is supported by BBSRC TRDF award BB/N023196/1, BBSRC National Capability Grants BB/J010375/1 and BB/CCG1720/1, BBSRC Institute Strategic Programme Grant BB/J004669/1 and BBSRC Core Strategic Programme Grant BB/J004669/1. This work was funded by a Wellcome Trust Investigator Award (100/974/C/13/Z), and the BBSRC Norwich Research Park Bioscience Doctoral Training Grant (BB/M011216/1, supervisor LJH, student CAG); Institute Strategic Programme Gut Microbes and Health BB/R012490/1, and Institute Strategic Programme Gut Health and Food Safety BB/J004529/1 to LJH. Isolation work was funded by a Microbiology Society Research Visit Grant to TCB. LH is in receipt of an MRC Intermediate Research Fellowship in Data Science (UK MED-BIO, grant number MR/L01632X/1). We are grateful for the assistance of the Genomics Pipelines team at EI, as well as the NBI Computing Infrastructure for Science team. We are also grateful to research nurse Karen Few for obtaining consent from parents and collecting samples. We thank Chris Bennett and Sasha Stanbridge of the EI Communications team for producing the accompanying video.

The following reagent was obtained through BEI Resources, NIAID, NIH as part of the Human Microbiome Project: Genomic DNA from Microbial Mock Community B (Staggered, High Concentration), v5.2H, for Whole Genome Shotgun Sequencing, HM-277D.

## AUTHORS CONTRIBUTIONS

LJH, MDC and RML designed the research; CAG, DH, MK, RML, SC, and TCB performed research; RML contributed new software tools; CAG, LH, LJH, MDC, PC, RML, SM and SC analysed data; and CAG, LJH, MDC and RML wrote the paper.

## ONLINE METHODS

### Mock Community

#### DNA

We used genomic DNA from a microbial mock community used in the Human Microbiome Project (HM-277D, BEI Resources, Manassas, VA). This mock community contains a mixture of 20 bacterial strains containing staggered RNA operon counts. Details of the strains present in the community are indicated in **Supplementary Table 1**.

#### Illumina sequencing of mock community

Illumina compatible amplification-free paired end libraries were constructed with inserts spanning from 600 bp to >1000 bp. A total of 600 ng of DNA was sheared in a 60 μl volume on a Covaris S2 (Covaris, Massachusetts, USA) for 1 cycle of 40 s with a duty cycle of 5%, cycles per burst of 200 and intensity of 3. Fragmented DNA was then end-repaired using the NEB End Repair Module (NEB, Hitchin, UK), size selected with a 0.58x Hi Prep bead clean-up (GC Biotech, Alphen aan den Rijn, The Netherlands) and followed by A tailing using the NEB A tailing module (NEB) and ligation of adapters using the NEB Blunt/TA Ligase Master Mix (NEB). Three 1x bead clean-ups were then undertaken to remove all traces of adapter dimers. Library quality control was performed by running an Agilent BioAnalyser High Sensitivity chip and quantified using the Kappa qPCR Illumina quantification kit. Based on the qPCR quantification libraries were loaded at 9 pM on an Illumina MiSeq and sequenced with 300 bp paired reads.

#### MinION sequencing of mock community

MinION 2D libraries were constructed targeting inserts >8 kbp. A total of 1 μg of DNA was fragmented in a 46 μl volume in a Covaris G-tube (Covaris, Massachusetts, USA) at 6,000 rpm in an Eppendorf centrifuge 5417. Sheared DNA was then subjected to a repair step using the NEB FFPE repair mix (NEB, Hitchin, UK) and purified with a 1x Hi Prep bead clean-up (GC Biotech, Alphen aan den Rijn, The Netherlands). A DNA control was added to the repaired DNA and then end-repaired and A-tailed using the NEBNext Ultra II End Repair and A-Tailing Module (NEB), purified with a 1x Hi Prep bead clean-up and then the AMX and HPA MinION adapters ligated using the NEB Blunt/TA Ligase Master Mix (NEB). An HP tether was then added and incubated for 10 min at room temperature followed by a further 10 min room temperature incubation with an equal volume of pre-washed MyOne C1 beads (Thermo Fisher, Cambridge, UK). The library bound beads were washed twice with bead binding buffer (ONT) before the final library eluted via a 10 min incubation at 37 ºC in the presence of the MinION Elution Buffer. The final library was then mixed with running buffer, fuel mix and nuclease free water and loaded onto an R7.3 flow cell per the manufacturer’s instructions and sequencing data collected for 48 h.

#### Mock community data analysis

MinION reads were basecalled using the Metrichor service and downloaded as FAST5 files. NanoOK v0.54^27^ was used to extract FASTA files, to align (via the LAST aligner^60^) against a reference database of the 20 genomes and to generate an analysis report (**Supplementary Note 1**). A QC of the Illumina data was carried out with FastQC (https://www.bioinformatics.babraham.ac.uk/projects/fastqc/) to ensure read quality was within expected bounds. This demonstrated a mean QC of 30 up to base 250. We subsampled a random set of 1,000,000 reads (subsample.pl script, https://github.com/richardmleggett/scripts) to represent the yield of a MiSeq nano flow cell and ran Trimmomatic^61^ to remove remaining adaptor content and apply a sliding window quality filter (size 4, mean quality greater than or equal to 15). Illumina and MinION reads were then BLASTed against the NCBI nt database and results were imported into MEGAN6^29^ for taxonomic analysis. The Pearson coefficient was computed in Excel using species abundance counts exported from MEGAN.

### Clinical samples

#### Ethical approval and preterm sample collection

The Ethics Committee of the Faculty of Medical and Health Sciences in the University of East Anglia (Norwich, UK) approved subject recruitment for this study. Protocol for faeces collection was laid out by the Norwich Research Park (NRP) Biorepository (Norwich, UK) and was in accordance with the terms of the Human Tissue Act 2004 (HTA), and approved with license number 11208 by the Human Tissue Authority. Infants admitted to the Neonatal Intensive Care Unit (NICU) of the Norfolk and Norwich University Hospital (NNUH, Norwich, UK) were recruited by doctors or nurses with informed and written consent obtained from parents. Oral probiotic supplementation provided to the infants in this study contained *Bifidobacterium bifidum* and *Lactobacillus acidophilus* (i.e. Infloran^®^, Desma Healthcare, Switzerland) strains with a daily dose of 2 × 10^9^ of each species. Collection of faecal samples was carried out by researchers and stored at −80 °C prior to DNA extraction.

#### DNA extraction of faeces samples from preterm infants

Bacterial DNA was extracted using the FastDNA Spin Kit for Soil (MP) following the manufacturer’s instructions but extending the bead-beating step to 1 min, and eluting the DNA with 55 °C DES. The starting faecal material used for the DNA extraction was between 100 and 150 mg. The purity and concentration of the DNA was assessed using a NanoDrop 2000c Spectrophotometer and Qubit^®^ 2.0 fluorometer. Samples with DNA concentrations higher than 25 ng/μl were considered acceptable.

#### MinION shotgun library preparation

MinION 2D libraries were constructed as outlined for the mock (see above) except that for the R9.4 flow cells the final library was mixed with running buffer containing fuel mix, library loading beads and nuclease free water and loaded onto the flow cell per the manufacturer’s instructions. MinION 1D libraries were prepared by incubating 200 ng of DNA for with 2.5 μl FRM 1 min at 30 °C then 1 min at 75 °C followed by the addition of 1 μl RAD and 0.2 μl NEB Blunt/TA Ligase Master Mix (NEB) and a room temperature incubation for 5 min. The final library was then mixed with running buffer containing fuel mix, library loading beads and nuclease free water and loaded onto the flow cell per the manufacturer’s instructions. Further details on Genomic Sequencing kits and samples used in this study can be found in **Table 1**.

#### Illumina HiSeq 2500 shotgun library preparation

Libraries for samples (P10N, P10R and P10V) were prepared using TruSeq Nano DNA Library Prep Kit according to the manufacturer’s instructions and sequenced using the HiSeq Illumina 2500 machine with 150 bp paired end reads. The library for P8 was prepared as for the Amplification Free library for the mock (see above) and run at 9 pM on an Illumina MiSeq with a 2× 250 bp read metric.

#### 16S rRNA gene library preparation and bioinformatics analysis

16S rRNA gene libraries were prepared using bacterial DNA normalised to 5 ng ml-1 for sample P10N and P10V. V4 region of the 16S rRNA gene was amplified using primers, 5’ AAT GAT ACG GCG ACC ACC GAG ATC TAC A and, 5’ CAA GCA GAA GAC GGC ATA CGA GAT AAC T. PCR conditions used for this amplification were: 1 cycle of 94°C 3 min and 25 cycles of 94°C for 45 s, 55°C for 15 s and 72°C for 30 s using a 96 well Thermal Cycler PCR machine. Sequencing of the 16S RNA gene libraries was performed on the Illumina MiSeq platform with 250 bp paired end reads.

Bioinformatic analysis started by processing raw reads through quality control using FASTX-Toolkit^62^ keeping a minimum quality threshold of 33 for at least 50% of the bases. Reads that passed the threshold were aligned against SILVA database (version: SILVA_119_SSURef_tax_silva)^63^ using BLASTn^64^ (version 2.2.25, max e-value 10e-3) separately for both pairs. After performing the BLASTn alignment, all output files were imported and annotated using the paired-end protocol of MEGAN.

#### Time series study for infant P10

Illumina and MinION sequencing data for samples P10N, P10V and P10R from infant P10 were studied. For the Illumina samples, we removed PCR duplicates (remove_pcr_duplicates.pl script from https://github.com/richardmleggett/scripts), ran Trimmomatic^52^ to remove adaptors and applied a sliding window quality filter (size 4, mean quality greater than or equal to 15) and then randomly sub-sampled 1 million reads (subsample.pl script from same location). These reads were used as the input to a BLASTn search (version 2.2.29, max e-value 10e-3) of NCBI’s nt database. For the Nanopore sequencing, we took only the reads classified as ‘pass’ reads (defined as 2D reads with a mean Q value >9) and performed no further pre-processing before running BLASTn. Using MEGAN6, we removed reads matching Homo sapiens (accounting for < 0.1% per sample) and performed taxonomic analysis.

#### Real-time diagnostic study using MinION and NanoOK RT

One sample from infant P8 was sequenced and both 1D and 2D Nanopore libraries were prepared using the SQK-LSK108 Ligation Sequencing Kit 1D, SQK-RAD002 Rapid Sequencing Kit 1D and SQK-LSK208 Ligation Sequencing Kit 2D. For samples P49A, P250G, P106I and P116I, SQK-LSK108 Ligation Sequencing Kit 1D was used. For sample P103M, SQK-RAD004 Rapid Sequencing Kit 1D was used. Libraries were sequenced on a mixture of R9.4 and R9.5 flowcells, as shown in Table 1. MinKNOW software was used to collect signal data. As described in the main text, an attempt was made to use the Metrichor base calling service for the P8 2D library but this failed, instead local basecalling through MinKNOW was used, although at the time only basecalled the template strand, so could not generate more accurate 2D consensus reads. MinKNOW was also used for local basecalling of the other 1D libraries.

To enable real-time analysis of MinION data, new functionality was added to NanoOK^29^. The new software, NanoOK RT, monitors a specified directory for basecalled sequence files as they are created by MinKNOW. For efficiency files are grouped into batches of 500, and each batch was BLASTn searched (version 2.2.29) against the NCBI nt database (downloaded April 2017) and the CARD database (v1.1.1, downloaded October 2016) of antibiotic resistance genes^65^. NanoOK RT also writes out command files for MEGAN, which allows more detailed analysis of community composition, either as the run proceeds, or on completion. NanoOK RT is available as an extension to NanoOK, selectable as a run-time option, from https://github.com/richardmleggett/NanoOK.

Another bioinformatic tool, NanoOK Reporter, was also developed for this project, and provides a graphical user interface to monitor the run and view summaries of community composition and antibiotic resistance genes identified. NanoOK Reporter uses a Lowest Common Ancestor (LCA) Algorithm to assign reads to the lowest possible taxonomy level consistent with all good BLAST matches. Results are displayed on a taxonomy tree, donut plot, or as a summary table showing most abundant matches. The tool allowed the user to browse through data in real time as batches were processed, or after all of the results were in using their timestamps to indicate when a result was first obtained. Summary data can also be exported as plain text files and these were subsequently used for later analysis. The LCA algorithm is only appropriate for species assignment; for multiple AMR hits, only the highest scoring hit is used for each individual read. ‘Walkout’ analysis can be initiated by clicking an icon and will produce a pie chart showing the species containing antibiotic resistance genes, as well as generating a text file giving per-read analysis. The walkout analysis proceeds by examining each read that has a good quality hit to an AMR gene to see if it also has an ‘independent’ hit to the nt (or bacterial alias) database. In our experiments, we defined independence as being a match the stretched at least 50 bases away from the AMR gene in either direction. NanoOK Reporter is available from https://github.com/richardmleggett/NanoOKReporter. Documentation for NanoOK Reporter as well as a tutorial utilising the data from this publication is available at https://nanook.readthedocs.io/en/latest/reporter.html.

#### Generation of AMR gene heat maps (Figures 4b, 4d, 7b and Supplementary Figures 7c, 7d, and 10b)

We opened the CARD results using NanoOK Reporter and used the option to save summary data as a plain text file. This saved a text file for the analysis at each time point (here batches of timestamped 500 reads) summarising the counts of resistance genes identified up to that point. We took the latest time point file that the heat map was to show (e.g. 6 hrs) and extracted a list of the ARO (Antibiotic Resistance Ontology) numbers from the ID column. Each unique ARO was manually assigned to its corresponding antibiotic resistance group following the classification given by CARD database. We wrote a script (gather_heatmap_data.pl, available at https://github.com/richardmleggett/bambi) to take the summary files, together with this mapping and to generate a final file summarising hits per group at each time point. An R script (plot_card_heatmap.R, same GitHub repository) takes this file and renders the heat map.

#### Statistical analysis on number of reads

Read counts at different stages of the bioinformatics analysis are provided in **Table 1**. For comparative analysis, sequences were subsampled down to the read count of the sample with the lowest number of reads.

#### Isolation and biochemical characterisation of P8 *Klebsiella pneumoniae* strains

An aliquot (100 mg) of faecal sample was homogenised in 1 mL TBT buffer (100 mM Tris/HCl, pH 8.0; 100 mM NaCl; 10 mM MgCl_2_•6H_2_O) by pipetting and plate mixing at 1500 rpm for 1 h. Homogenates were serially diluted to 10^−4^ in TBT buffer. Aliquots of 50 μl were spread on MacConkey (Oxoid) agar plates in triplicate and incubated aerobically at 37 °C overnight.

Colonies were selectively screened for lactose-positive (i.e. pink) colonies. One colony of each morphology type was re-streaked on MacConkey agar three times to purify. Biochemical characterisation was performed using API 20E tests (Biomerieux) according to manufacturer’s instructions.

#### 16S rRNA phylogenetic analysis of P8 *Klebsiella pneumoniae* isolates

Sequences of the 16S rRNA gene from nine *Klebsiella pneumoniae* isolates were prepared to perform the phylogenetic analysis. We extracted DNA using FastDNA Spin Kit for Soil (MP) following the manufacturer’s instructions and then amplified the 16S rRNA gene employing Veriti™ 96 well Thermal Cycler (Applied Biosystems, USA), master mix from Kapa2G Robust PCR reagents (Kapa Biosystems, USA) and primers: fD1 (5’-AGA GTT TGA TCC TGG CTC AG -3’), fD2 (5’-AGA GTT TGA TCA TGG CTC AG -3’) and rP1 (5’ -ACG GTT ACC TTG TTA CGA CTT -3’)^66^. PCR amplification conditions were: 1 cycle of 94 °C for 5 min, followed by 35 cycles of 94 °C for 1 min, 43 °C for 1 min, and 72 °C for 2 min followed by a final strand extension at 72 °C for 7 min. Amplicons were sequenced using automated Sanger sequencing service (Eurofins Genomics, Luxembourg).

Partial 16S ribosomal RNA sequences (~900 positions) of nine isolates of *Klebsiella pneumoniae*, obtained using automated Sanger sequencing service (Eurofins Genomics, Luxembourg), were compared for similarity. Multiple sequence alignments were performed with SINA (SILVA Incremental Aligner) (Pruesse et al., 2012) (https://www.arb-silva.de/aligner/), and manually curated for quality. The nucleotides were coloured using Boxshade version 3.21 (http://www.ch.embnet.org/software/BOX_form.html). The similarity/identity matrix between the sequences was calculated using MatGAT, version 2.01 (Matrix Global Alignment Tool) with BLOSUM 50 alignment matrix^67^.

#### Determination of Minimal Inhibitory Concentration (MIC) for P8 *Klebsiella pneumoniae* isolate

Calculation of the antibiotic Minimal Inhibitory Concentration (MIC) of growth of a *Klebsiella pneumoniae* isolate from baby P8 (sample P8E) was performed using the broth microdilution method^68^. Serial two-fold dilutions of the most common antibiotics used at NICU (benzylpenicillin, gentamicin, vancomycin, metronidazole, meropenem and cefotaxime) were added to sterile nutrient broth. Bacterial inoculum of the isolate was prepared using 10 μl from a fresh overnight culture, and tests were done in triplicate. Microplates were incubated for 24 h at 37 °C under aerobic conditions. Cell density was monitored using a plate reader (BMG Labtech, UK) at 595 nm. MICs were determined as the lowest concentration of antibiotic inhibiting any bacterial growth.

#### DNA extraction from P8 *Klebsiella pneumoniae* isolate for WGS analysis

An overnight (10 ml) culture of the isolate was centrifuged at 4000 rpm for 10 min, re-suspended in 30 ml of Phosphate Buffered Saline (PBS) (Sigma-Aldrich, UK) and centrifuged again. The pellet was then re-suspended in 2 ml of 25% sucrose (Fisher Scientific, USA) in TE (10 mM Tris (Fisher Scientific, USA) and 1 mM EDTA at pH 8.0 (VWR Chemicals, USA); 50 μl of Roche Lysozyme (Roche Molecular Systems, UK) at 100 mg/ml in 0.25 M Tris pH 8.0 was added. The mixture was incubated at 37 °C for 1 h, and 100 μl of Proteinase K at 20 mg/ml (Roche Molecular Systems, UK), 30 μl of RNase A at 10 mg/ml (Roche Molecular Systems, UK), 400 μl of 0.5 M EDTA pH 8.0 (VWR Chemicals, USA), and 250 μl of freshly prepared 10% Sarkosyl NL30 (Sigma-Aldrich, UK) were added. The mixture was then incubated on ice for 2 h and subsequently transferred to a water bath at 50 °C overnight. Next, E Buffer (10 mM Tris pH8.0 (Fisher Scientific, UK)) was added to the sample to a final volume of 5 ml, mixed with 5 ml Phenol:Chloroform:Isoamyl Alcohol (PCIA) (25:24:1) (Sigma-Aldrich, UK) in a Qiagen MaXtract High Density tube (Qiagen, DE) and centrifuged for 15 min at 4000 rpm. The aqueous phase was transferred into a new Qiagen MaXtract High Density tube, made up with E Buffer to the volume of 5 ml if necessary, mixed with 5 ml of PCIA, and centrifuged for 10 min at 4000 rpm. This procedure was repeated, with 5 min centrifugation time. Next, the aqueous phase was transferred into a Qiagen MaXtract High Density tube, made up to 5 ml with E Buffer if necessary, mixed with 5 ml of Chloroform:Isoamyl Alcohol (CIA) (24:1) (Sigma-Aldrich, UK), and centrifuged for 5 min at 4000 rpm. The CIA step was repeated once more, after which the final aqueous phase was transferred into a sterile Corning TM 50ml centrifuge tube, and 2.5 volumes of ethanol (Ethanol absolute AnalaR NORMAPUR^®^, VWR Chemicals, USA) were added. The sample was incubated for 15 min at −20 °C, then centrifuged for 10 min at 4000 rpm and 4 °C. Finally, the DNA pellet was washed with 10 ml of 70% ethanol and centrifuged at 4000 rpm for 10 min twice, dried overnight, and re-suspended in 300 μl of E Buffer.

#### Whole genome sequencing library preparation and sequencing of P8 *Klebsiella pneumoniae* isolate

DNA samples containing 500 ng genomic DNA were analysed. DNA was sheared into fragments of 400-600 bp using a Covaris plate with glass wells and AFA fibres. SPRI clean-up was used to remove smaller sized fragments and concentrate the sheared DNA samples. Whole genome library construction performed by a liquid handling robot comprised end repair, A-tailing and adapter ligation reactions. Adapter ligated samples were subsequently amplified using the following PCR conditions: 5 min 95°C, 10 cycles of (30 sec 98°C, 30 sec 65°C, 1 min 72°C) and 10min at 72°C. LabChip GX was then used to size and assess the quality of the libraries and determine pooling volumes for each library using Beckman Coulter Biomek NXp (span-8). Libraries were prepared using Agilent Technologies Sure Select Custom Library Prep kit. Final pools were finally loaded on the HiSeq 2500 sequencers.

#### Assembly and detection of AMR genes from whole genome sequencing isolate (P8*Klebsiella pneumoniae*) and MinION metagenomic samples (P8, P10N, P10R and P10V)

Presence or absence of AMR genes was performed on one *Klebsiella pneumoniae* isolated from sample P8 benchmarking two different sequencing platforms: MinION and Illumina HiSeq 2500. Sequencing data from the MinION run was assembled using Canu version^69^ (v1.5) corrected with Racon^70^ (v1.3.1) and polished with nanopolish^71^ (v0.9.0). Sequencing data from the Illumina HiSeq 2500 run was assembled using Velvet. Contigs were BLASTn searched (v2.2.29) against the CARD database (v1.1.1) with maximum e-value 10e^−3^. Further filtering of (90% identity, 90% coverage) and (80% identity, 80% coverage) were carried out to produce Supplementary Figure 5.

#### Bifidobacterium bifidum assembly

The P103M MinION reads were aligned against the *Bifidobacterium bifidum* PRL2010 reference sequence from NCBI using minimap2. All mapping reads with an alignment quality of 10 or greater were used as the input to the assembly. Reads were processed with porechop to remove adaptors, prior to assembly with Canu (v1.5) and polishing with nanopolish (v0.9.0). The output contigs from this step were then used as the input to minimus2, resulting in 3 final contigs. Accuracy of assembly was assessed using dnadiff, part of MUMmer^72^ and with BRIG^73^.

